# Selective Targeting of Ras Mutant Cancers via a New Small Molecule

**DOI:** 10.1101/2020.12.23.423300

**Authors:** Bhairavi Tolani, Anna Celli, Yanmin Yao, Yong Zi Tan, Richard Fetter, Christina R. Liem, Adam J. de Smith, Thamiya Vasanthakumar, Paola Bisignano, Ian B. Seiple, John L. Rubinstetin, Marco Jost, Jonathan S. Weissman

## Abstract

Mutations in the Ras family of oncogenes are implicated in 33% of human cancers, making Ras an intensely pursued target in drug discovery. As an alternative to direct pharmacological inhibition of Ras, we looked for sensitivities in RAS mutant cells. Using a small molecule screen in cell lines with mutations in Ras and its effector Raf, we discovered 249C as a Ras-mutant selective cytotoxic agent against a spectrum of RAS-mutant cancers. By combining CRISPR chemical-genetic screening, comparative profiling and chemoproteomics, we identified that 249C binds to a unique subunit on vacuolar (V)-ATPase with nanomolar affinity, inhibiting its biochemical activity and, unexpectedly, altering V-ATPase translocation in Ras-induced macropinocytosis. Via binding to V-ATPase, 249C prevents lysosomal acidification and inhibits autophagy and macropinocytosis pathways that several Ras-driven cancers rely on for survival. In characterizing 249C’s mechanism, we show that potency varies with the identity of the RAS driver mutation highlighting a mutant-specific dependence on autophagy and macropinocytosis. Indeed, 249C potently inhibits tumor growth without adverse side effects in a mouse xenograft model of *KRAS*-driven non-small cell lung cancer. These data establish proof-of-concept for targeting V-ATPase as a way to indirectly target specific Ras mutants, and provide a fundamental link between V-ATPase localization and specific Ras mutant tumor-related activity.

## INTRODUCTION

Mutations that constitutively activate members of the Ras family of oncogenes (H-Ras, N-Ras, and K-Ras) are collectively responsible for about one third of all human cancers; *KRAS* mutations in particular are implicated in the most fatal malignancies: pancreatic (91%), colon (42%) and lung (33%)^1–4^. Despite decades-long efforts, we still lack clinically-approved drugs for the majority of oncogenic *KRAS* variants (as we await clinical data from Amgen’s *KRAS*G12C trials), let alone a compound with pan-*KRAS* activity, in large part due to substantial challenges in directly inhibiting mutant *KRAS*. As an alternative strategy for pharmacological intervention, efforts to map potentially druggable neomorphic dependencies of *KRAS* mutant cancers have identified, among others, metabolic adaptations concomitant with activating *RAS* mutations, including high levels of basal autophagy, a phenomenon termed “autophagy addiction”^5–9^, and constitutive activation of macropinocytosis (MP)^10^. Both autophagy and MP provide nutrients to promote tumor growth by degrading macromolecules in acidic lysosomes produced by vacuolar H^+^ ATPases (V-ATPases)^11,12^, and V-ATPase has been reported as an essential regulator of *RAS*-induced MP for nutrient supply, suggesting that targeting V-ATPase could be exploited to curtail the metabolic adaptation of *RAS*-mutant cells^10^.

V-ATPase is a multi-subunit transmembrane complex that operates as an ATP-driven rotary proton pump, coupling ATP hydrolysis in its peripheral V_1_ domain to proton translocation from the cytoplasm to the lumen of organelles through its membrane integral Vo domain. Through this activity, V-ATPase is essential for lysosome acidification and thus lysosomal degradation^13,14^ and more broadly regulates a wide array of membrane trafficking and intracellular transport processes^15^. Although specific V-ATPase inhibitors such as Bafilomycin A1 (BafA1) have been developed and serve as important tool compounds, their clinical use is limited by toxicity resulting from acidosis and hypoxia^16^. Additional autophagy-modulating agents have been described, such as the repurposed anti-malarial compound chloroquine (CQ) and its less toxic derivative hydroxychloroquine (HCQ), which impair lysosomal acidification by targeting Palmitoyl-protein thioesterase 1 (PPT1), but these compounds have severe side effects by interfering with human *ether-à-go-go*-related gene (hERG) responsible for electrical activity of the heart^17,18^. In over 60 clinical trials (completed or ongoing), the use of CQ or HCQ alone as a monotherapy do not demonstrate significant therapeutic efficacy^19^. Although second-generation analogs of HCQ such as ROC325, Lys05, DC661, and DQ661 show enhanced preclinical lysosomal-autophagic inhibition and antitumor activity as single agents *in vitro* and *in vivo*, it remains to be determined whether any of them will show efficacy in clinical investigations while avoiding unacceptable toxicities^17,20^. These limitations motivate an urgent need to identify and develop additional autophagy/MP inhibitors with clean safety profiles.

Here, we report the discovery and characterization of a class compounds with potency in killing *KRAS* mutant cells and good pharmacological properties. By combining chemoproteomics, comparative profiling, and chemo-genetic CRISPR screening, we determined that our lead compound 249C inhibits autophagy/MP through binding to V-ATPase, resulting in the inactivation of lysosomal degradation and selective killing of *KRAS/BRAF* mutant cells.

## RESULTS

### Cell Proliferation Screen Reveals 249C as Potent Inhibitor of Ras and Raf Mutant Cancer Cells

To discover new lead compounds for *Ras* and *Raf* mutant cells, we synthesized and screened ~300 rationally designed small molecules for anti-proliferative activity against three human cancer cell lines with mutations in *Ras* or *Raf*: A549 (*KRAS*G12S), LOX-IMVI (*BRAF*V600E), and MelJuso (*HRAS*G13D and *NRAS*Q61L). Using structure-activity relationships, we observed that several compounds potently reduced viability of all 3 cell lines (**Fig. 1a**). Simultaneous evaluation of Lipinski’s drug-like properties and half-maximal inhibitory concentration (IC_50_) values revealed 249C as the lead compound (**Fig. 1b**). To further evaluate determinants of sensitivity to 249C, we measured its potency against 53 cell lines (lung, pancreatic, breast, ovarian, colorectal/gastric, prostate, liver/hematological, glioma and melanoma) (**Fig. 1c**). Comparing IC_50_’s to gene mutation data (CCLE) revealed that the presence of at least one of *KRAS*, *HRAS*, *NRAS*, and/or *BRAF* mutations was significantly associated with lower IC_50_ values (*p*=1.24 × 10^−6^), with median IC_50_ values of 65 nM in *KRAS*/*HRAS*/*NRAS*/*BRAF* mutant cells versus 365 nM in cells with no mutations in these genes (**Fig. 1d**). We next mapped the response of A549 cells to these compounds using mass spectrometry (LC–MS/MS)-based protein profiling coupled with quantitative tandem mass tags labeling^21^. Upon treatment of cells with vehicle or molecules 68, 226C and 249C, we observed substantially elevated levels of the autophagy receptor SQSTM1/p62 common to this family of compounds (**Fig. 1e**) in a time-dependent fashion (**Fig. 1f**), suggesting that these compounds impact autophagy.

**Figure 1:**
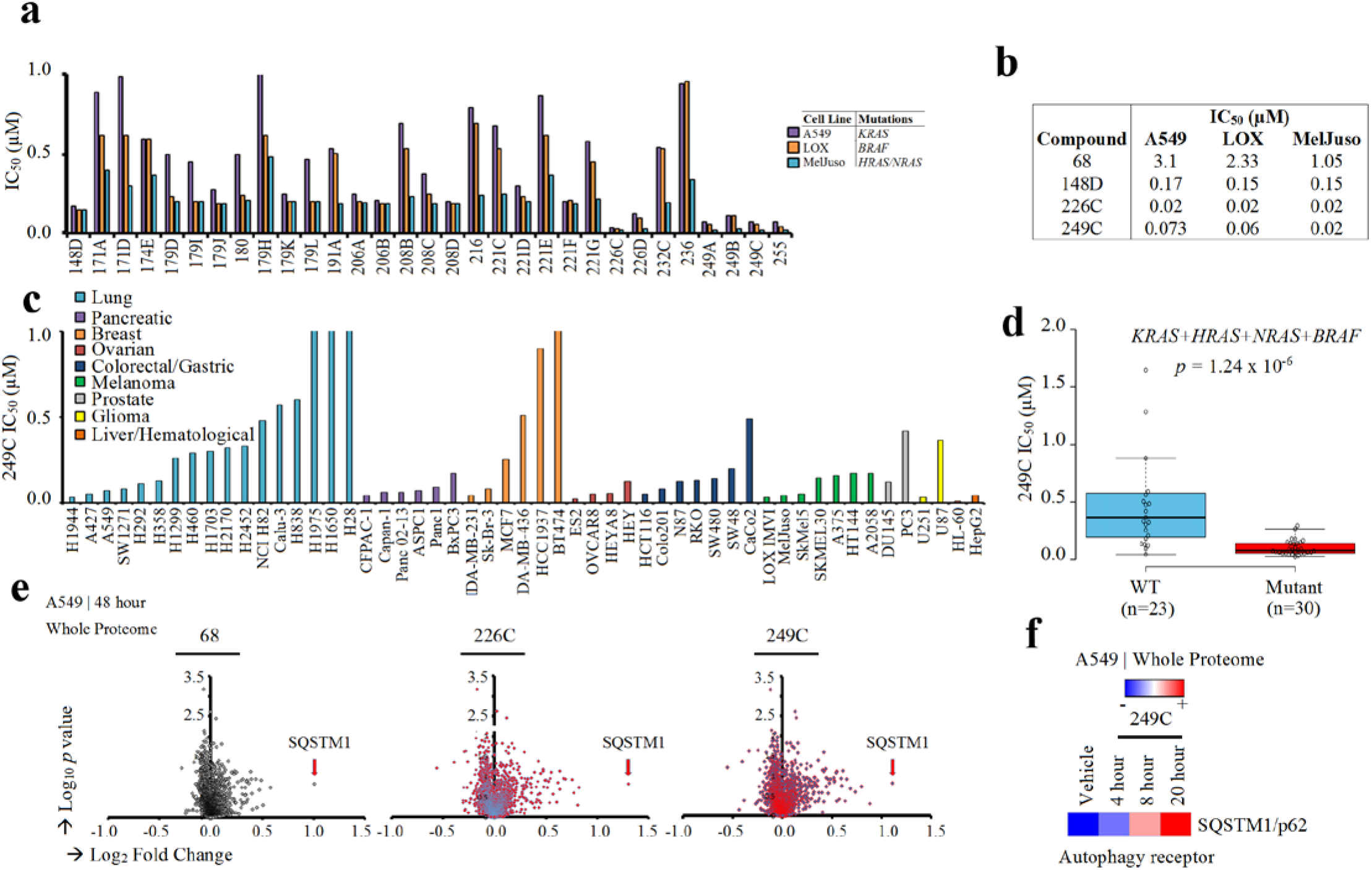
Optimization of Potency and Drug Properties of Small Molecule Inhibitors of *Ras-* and *Raf-* mutant Cells using Viability Screens. (**A**) Cell viability screen of ~300 small molecules in A549 (*KRAS*G12S), LOX IMVI (*BRAF*V600E), and MelJuso (*HRAS*G13D and *NRAS*Q61L). Cells were treated with vehicle DMSO or increasing concentrations of small molecules in triplicate for 72 h and ATP content/viability was measured using CellTiter-Glo Luminescent Cell Viability Assay. The results shown for all cell viability figures are representative of at least two independent experiments. The top 30 compounds (p<0.05 compared to vehicle-treated controls) with the lowest IC_50s_ are presented. (**B**) Evolution of the Structure Activity Relationship (SAR) and IC_50_s of the first (68) and last (249C | lead due to drug-like properties) compounds in the screen. (**C**) 249C sensitivity for a panel of 53 cancer cell lines (including lung, pancreatic, breast, ovarian, hematological/liver, prostate, colorectal cancers, glioma and melanoma). (**D**) IC_50_ values stratified by the mutation status of *KRAS, HRAS, NRAS,* and *BRAF* (Wilcoxon rank sum test | *p* = 1.24 × 10^−6^). (**E**) Quantitative whole proteome analysis of the effects of small molecule treatment by mass spectrometry. A549 cells were treated with DMSO or 68, 226C or 249C for 48 h prior to analysis of proteomes with TMT labeling followed by quantification relative to the DMSO control. Autophagy receptor, SQSTM1, was identified as highly up-regulated and common for our family of compounds by proteomic mass spectrometric screening in small molecule-treated A549 mutant *KRAS* cells. (**F**) Quantitative whole proteome analysis of 249C-treated A549 cells over time relative to DMSO control.

### 249C Inhibits Autophagy and Phenocopies the V-ATPase Inhibitor Bafilomycin A1

To characterize the mode of action of our lead compound 249C, we used comparative profiling in a panel of primary cell-based systems designed to model disease biology^22^. Compared to 4,000 other drugs, the phenotypes elicited by 249C most closely resembled those elicited by the V-ATPase inhibitor BafA1 (Pearson *r* = 0.95 across all measurements, (**Fig. 2a, 2b**). 249C also exhibited similarity, albeit to a lesser degree, to the mTOR inhibitors Rapamycin and Torin1 and the antimalarials CQ and quinacrine. 249C and BafA1 elicited progressive accumulation of SQSTM1/p62 and LC3-II in A549 cells, consistent with a block in autophagic flux; by contrast, the autophagy inducer Rapamycin did not (**Fig. 2c**). Using transmission electron microscopy, we further observed late-stage autophagy inhibition and increased autophagic vesicle accumulation in A549 cells treated with 249C and BafA1 (**Fig. 2d-j**). Finally, treatment with both 249C and BafA1 caused a reduction in the presence of acidic organelles (red: **Fig. 2k** – top panel) and an increase in cellular pH (yellow: acidic; blue: neutral | **Fig. 2k** – bottom panel) of A549 cells relative to DMSO and Torin1, confirming that 249C blocks lysosomal acidification and autophagy progression.

**Figure 2.**
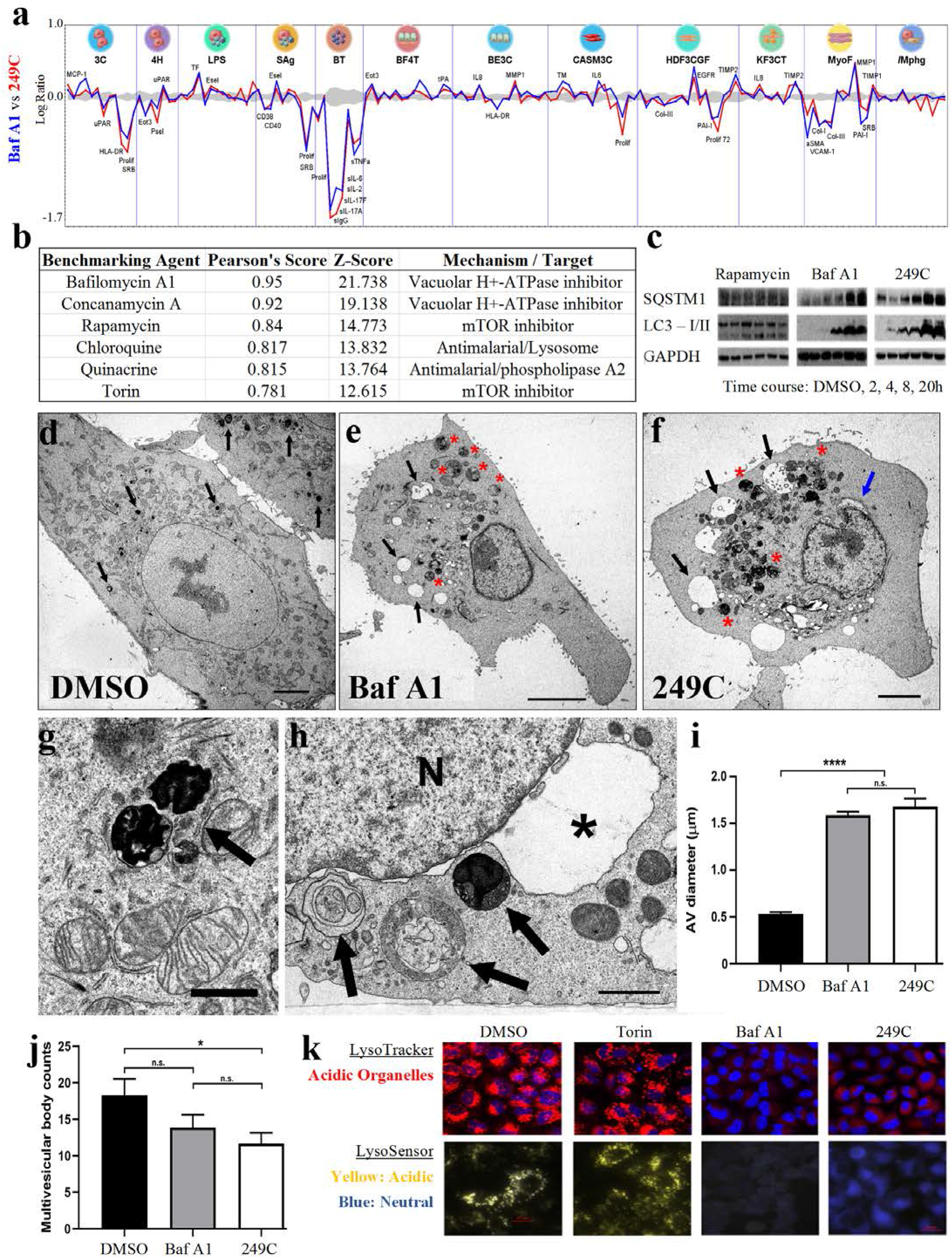
Lead Small Molecule 249C Phenocopies V-ATPase Inhibitor BafA1. (**A**) Overlay of 148 biomarker responses in a panel of 12 primary cell-based systems for **249C** and **BafA1**. BafA1 had the highest similarity out of 4000 molecules. (**B**) Pearson correlation and *z*-scores for observed phenotypes of 249C compared to known database drugs. (**C**) Treatment with 249C or the autophagy inhibitor BafA1 resulted in upregulation of autophagy markers SQSTM1/p62 and LC3-I/II over time but not with autophagy inducer, Rapamycin. (**D**) Representative electron micrographs of A549 cells treated with the DMSO vehicle, BafA1 (**E**) or 249C (**F**). Arrows in (D): very densely staining phagosomes/lysosomes. (**G**) shows two of these structures at higher magnification from a DMSO treated cell (arrow). Arrows in (E) and (F): large, clear vacuoles within the cells that are absent in (A); red asterisks in (E) and (F) large multivesicular autophagic vesicles (AVs). (**H**) Multivesicular AVs in a cell treated with 249C (arrows). The nucleus (N) in this cell exhibits a greatly distended portion of the nuclear envelope (asterisk) that resembles the large vacuoles seen in the cytoplasm of Baf A1- and 249C-treated cells. (**I**) Quantification of mean + SEM of the size of the multivesicular AVs and number of multivesicular bodies (**J**) from counts of DMSO -,Baf A1- and 249C-treated cells. Scale bars in (D-F), 5 μm; (G-H), 1 μ m. * p<0.05; **** p<0.0001; n.s. not significant. (**K**) Treatment with autophagy inhibitor, BafA1, and 249C decreased staining of acidic organelles (red | LysoTracker) and resulted in an increase in pH (yellow: acidic; blue: neutral | LysoSensor) relative to the DMSO control and autophagy inducer, Torin1, in A549 *KRAS* mutant cells. Representative images from n >3.

### Chemical-genetic Screen Implicates V-ATPase as the Cellular Target of 249C

To further pinpoint the molecular target of 249C, we turned to CRISPR interference (CRISPRi)-mediated chemical-genetic screening, which has emerged as a robust strategy for small molecule target identification^23^. We conducted these screens in breast adenocarcinoma MDA-MB-231 (*KRAS*-G13D and *BRAF*-G464V) cells, which are sensitive to 249C treatment. Briefly, we infected MDA-MB-231 cells stably expressing the dCas9-KRAB CRISPRi effector^24^ with the genome-scale hCRISPRi-v2 single-guide RNA (sgRNA) library (targeting 18,905 genes)^25^, harvested a subpopulation at the outset of the experiment (t_0_), and then cultured the remaining cells without (DMSO) or with 249C treatment. Finally, we used next-generation sequencing to determine how each sgRNA impacted sensitivity to 249C (ρ) and growth in the absence of 249C (γ) (**Fig. 3a**)^26^. The resulting untreated growth phenotypes were well-correlated with those from other cell types, suggesting that our screen faithfully captured phenotypes resulting from gene knockdown. Among the 147 genes for which knockdown strongly affected 249C sensitivity (**Fig. 3b**), we were intrigued to find 5 V-ATPase subunits (*ATP6V1H, ATP6V1A, ATP6V0A2, ATP6V0A3/TCIRG1*, and A*TP6V0E1*), because 249C essentially phenocopied the V-ATPase inhibitor BafA1 in our previous assays (Fig. 2c). Indeed, knockdown of several V-ATPase subunits sensitized cells to 249C; the regulatory H subunit ATP6V1H in particular was one of the most strongly sensitizing hits. By contrast, knockdown of the poorly characterized subunits *ATP6V0E1* and *ATP6V0A3* protected cells against 249C. Individual re-tests using internally controlled drug sensitivity assays further confirmed that knockdown of multiple V-ATPase subunits sensitized MDA-MB231 cells to 249C and BafA1 (target: ATP6VoC)^27^ (**Fig. 3c**). Thus, genetic depletion of V-ATPase sensitizes cells to 249C, further suggesting that 249C’s cytotoxicity arises from inhibition of V-ATPase activity.

**Figure 3.**
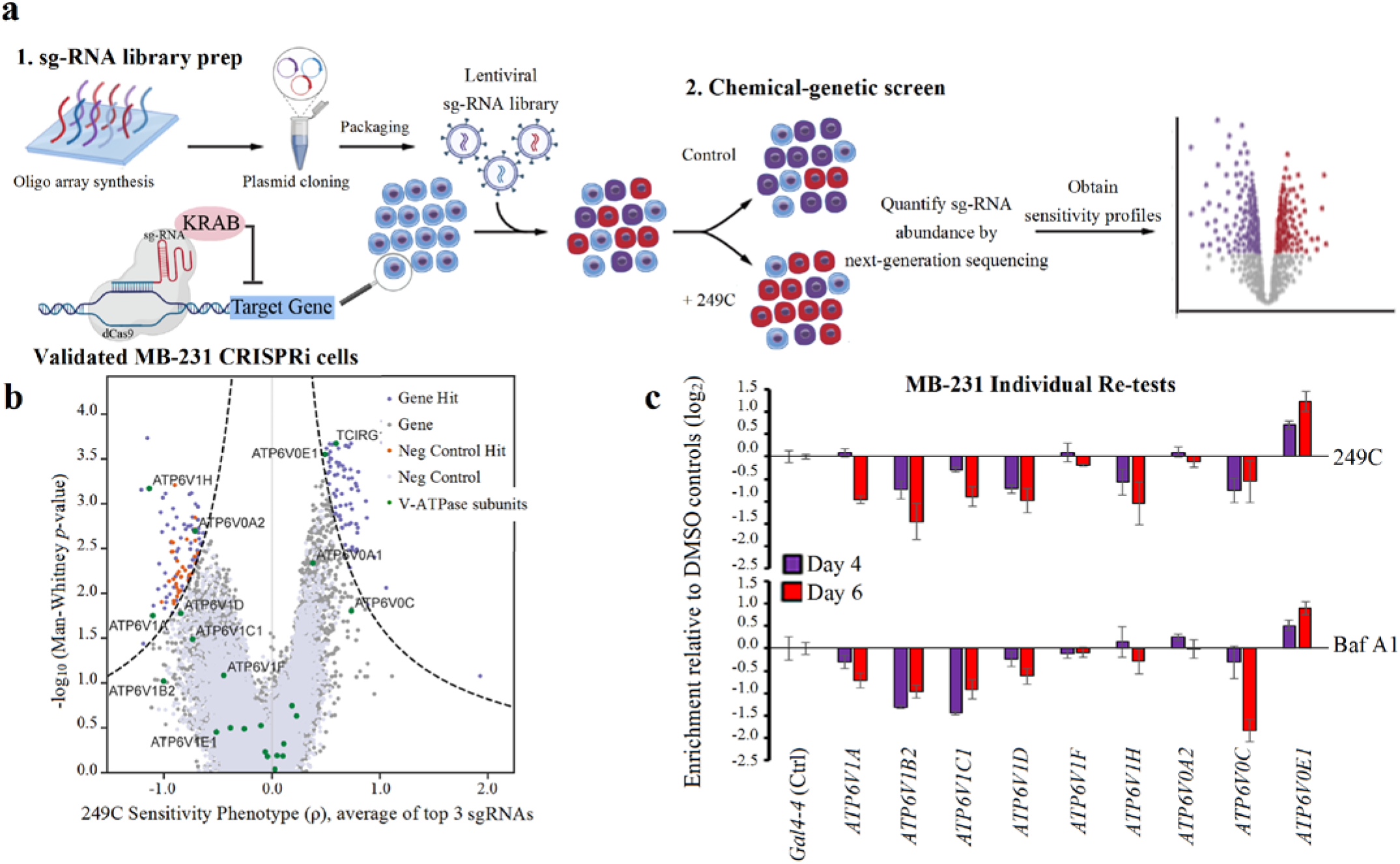
Genome-scale Chemical-genetic CRISPRi Screen Implicates V-ATPase as 249C’s Molecular Target. (**A**) Schematic illustration of the genome-wide CRISPRi chemical-genetic screen. (**B**) Volcano plot of 249C sensitivity phenotype from genome-scale CRISPRi screen. Phenotypes from sgRNAs targeting the same gene were collapsed into a single sensitivity phenotype for each gene using the average of the top three scoring sgRNAs and assigned a *p*-value using the Mann-Whitney test of all sgRNAs targeting the same gene compared to the non-targeting controls. Negative control genes were randomly generated from the set of non-targeting sgRNAs. Dashed line represents discriminant score ≥7, calculated as phenotype *z*-score ∙ −log_10_(*p*-value), with the *z*-score defined from the standard deviation of the negative control genes. V-ATPase genes are shown in green. (**C**) Internally controlled individual re-tests for 249C and Baf A1 sensitivity assays performed with sgRNAs targeting (*ATP6V1 –A, B2, C1, D, F, H* and *ATP6V0 – A2, C, E1*) or a non-targeting control sg-RNA (*Gal4-4*) in MDA-MB-231 CRISPRi cells. Cells transduced with the sg-RNA expression constructs (marked with blue fluorescent protein [BFP]) were left untreated or treated with 249C/Baf A1 4 days after transduction. Enrichment of sg-RNA-expressing cells was measured after treatment on the days indicated by flow cytometry as the enrichment of BFP (n = 3). Data presented: mean ± SD for replicate infections and treatments (n = 3). (Created with BioRender)

### 249C Deregulates V-ATPase Assembly, Inhibits Mammalian V-ATPase Activity and Binds ATP6V1H

With multiple independent assays pointing toward V-ATPase as the target of 249C, we proceeded to validate the biochemical and biophysical effects of 249C on V-ATPase activity. V-ATPase activity depends upon reversible assembly of the V1 and Vo domains and is nutrient-dependent; increased V1-Vo assembly increases catalytic activity^15^. To test whether 249C alters localiztion of V-ATPase subunits, we evaluated the partitioning of V-ATPase subunits into the cytosolic [containing the V_1_ domain composed of eight subunits (A–H) responsible for ATP hydrolysis] and membrane fractions [membrane-embedded Vo domain comprised of six subunits (a, c, c′ (absent in mammals), c′′, d and e) responsible for proton translocation]^15^ of 249C-treated HEK293T cells. Treatment with 249C but not BafA1 led to an increase in membrane-associated subunit B2 (cytosolic V_1_ domain), possibly indicating increased V-ATPase assembly, perhaps as part of its unique mechanism (**Fig 4a-b**). To independently corroborate the effect of 249C on V-ATPase and to probe whether proton pumping is impacted in the complexes on lysosomes, we measured lysosome acidification using a pH-dependent fluorescent probe, FITC-Dextran. Both 249C and BafA1 inhibited V-ATPase-dependent proton transport in mammalian cells, as measured by ATP-dependent fluorescence quenching of FITC-loaded lysosomes (**Fig. 4c**)^28^. Proton translocation through the Vo region is powered by ATP hydrolysis in the V_1_ region. In addition to blocking acidification, both 249C and BafA1 blocked ATP hydrolysis of intact mammalian V-ATPase purified from pig kidneys at concentrations as low as 1 μM, as measured by a standard biochemical V-ATPase activity assay (**Fig. dc**). To directly measure binding of 249C to V-ATPase, we performed bio-layer inferometry (BLI) with both the purified intact mammalian V-ATPase complex and the recombinant ATP6V_1_H subunit using biotinylated-249C loaded onto streptavidin sensors, which revealed binding with nanomolar affinities (*K_ds_*: complex = 23 nM; H sub = 501 nM) (**Fig. 4e-f**). Altogether, these results establish that 249C directly binds to and inhibits mammalian V-ATPase.

**Figure 4:**
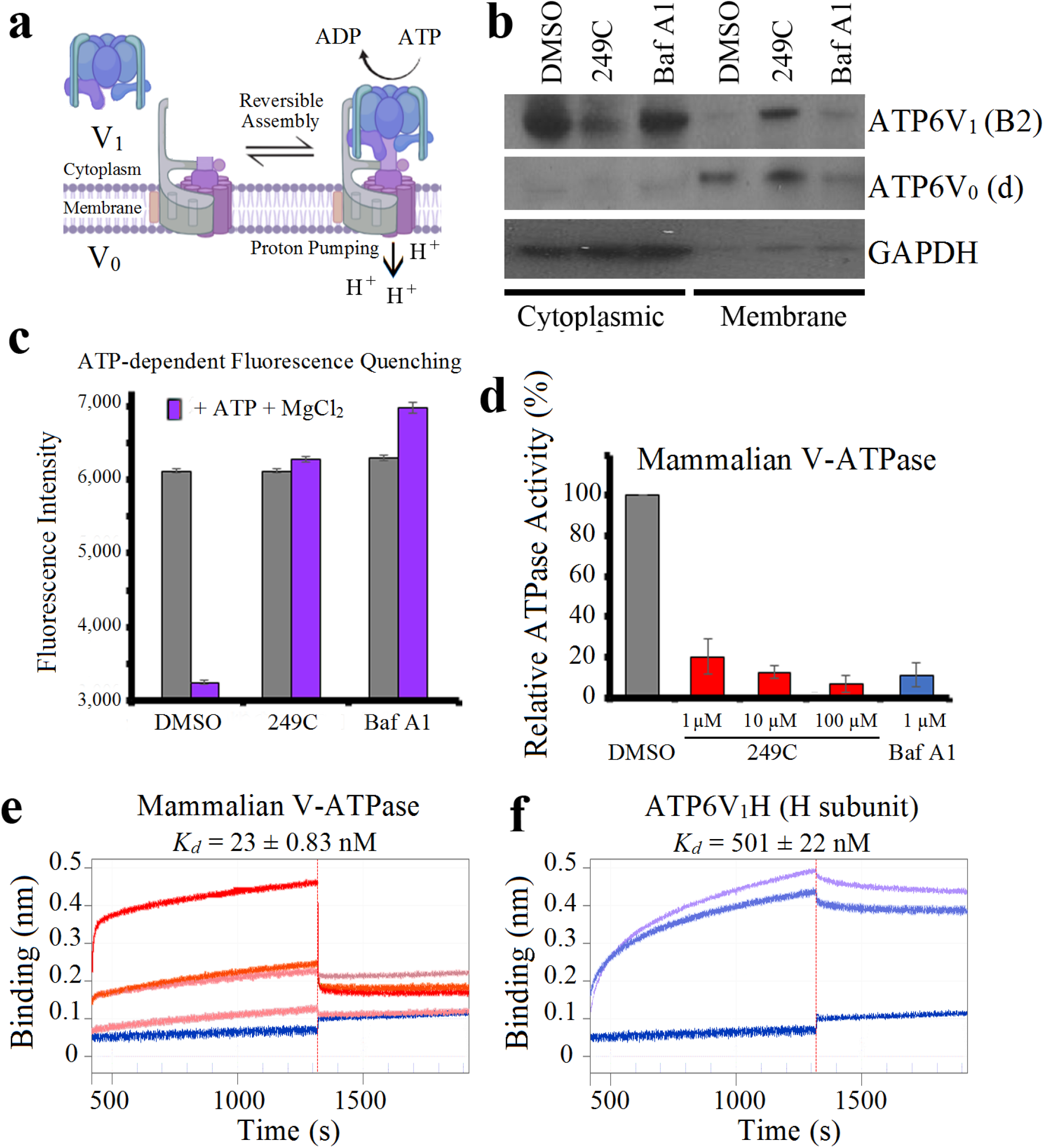
Validation of Functional Biochemical and Biophysical Effects of 249C on V-ATPase Activity. (**A**) Model of the reversible assembly of V-ATPase. Cytosolic V_1_ subunits are depicted in *blue* and membrane V_o_ subunits in *purple*. Upon V-ATPase assembly, ATP is hydrolyzed to ADP accompanied by proton (H^+^) pumping and luminal acidification (decrease in pH) (Created with BioRender). (**B**) HEK293T cells were treated with 249C or BafA1 and cell homogenates were prepared, separated into membrane and cytosolic fractions and analyzed by Western blotting using antibodies against subunit B2 as a measure of V1, subunit Vod as a loading control for the membrane fraction, and GAPDH as a loading control for the cytosolic fraction (see Methods). The amount of subunit B2 present in the membrane fraction indicates the amount of assembled V-ATPase. A representative Western blot is presented. (**C**) After 1 h of 249C or Baf A1 treatment, HEK293T cells were allowed to take up FITC-Dextran by endocytosis, and the dye was chased to the lysosomal compartment (see Methods). Cells were mechanically lysed, and a fraction containing FITC-Dextran-loaded lysosomes was isolated by sedimentation centtrifugation. Fluorescence was measured over time to assess pH-dependent quenching following addition of 1 mM magnesium-ATP (predetermined). ATP-dependent fluorescence quenching was not observed for 249C- or Baf A1- treated samples. Representative of 5 individual experiments with n = 2 for each is presented. (**D**) Mammalian V-ATPase activity measured in the absence and presence of 1 μM 249C and 1 μM Baf A1. Data shown are mean ± SD; n = 3. Bio-layer inferometry (BLI) of (**E**) the entire mammalian V-ATPase complex (30, 60, 120, 240 nM) and (**F**) the individual human H subunit (ATP6V1H) (16,800 and 21,000 nM) against biotinylated-249C loaded onto streptavidin sensor tips. V-ATPase complex: *K_d_* = 23 ± 0.83 nM; H subunit: *K_d_* = 501 ± 22 nM. Representative of two independent experiments.

### 249C Selectively Inhibits Ras/Raf-Induced Macropinocytosis and Alters ATP6V1A Cellular Localization

The most frequent *KRAS* mutations across all human cancers occur at codons 12 or 13 with replacement of glycine to other amino acids: G12D (35%), G12V (24%), G13D (13%)^29^. To test the impact of 249C on cells carrying individual Ras/Raf mutations, we measured the potency of 249C against immortalized mouse embryonic fibroblasts (MEFs) genetically engineered to be “Ras-less” with subsequent stable transduction of WT or mutant *KRAS* isoforms^30^ obtained from the NCI. Strikingly, 249C exhibited increased potency against cells expressing *KRAS/BRAF* mutants rather than the WT genes (**Fig. 5a**). Moreover, potency depended on the identity of the mutant [IC_50_ values: WT (1.25 μM), G13D (0.07 μM), G12V (0.15 μM), G12S (0.23 μM), G12D (0.3 μM), Q61L (0.31 μM), G12C (0.44 μM), Q61R (0.55 μM), *BRAF*V600E (0.11 μM)], suggesting that susceptibility to V-ATPase inhibition differs across *KRAS*/*BRAF* mutants (**Fig. 5a**). Treatment with BafA1 showed similar profiles except for G12C, G12D, and Q61L, whereas the HCQ analog (DC661) showed substantial differences and the *KRAS*G12C inhibitor (AMG510) specifically killed cells expressing *KRAS*G12C.

**Figure 5:**
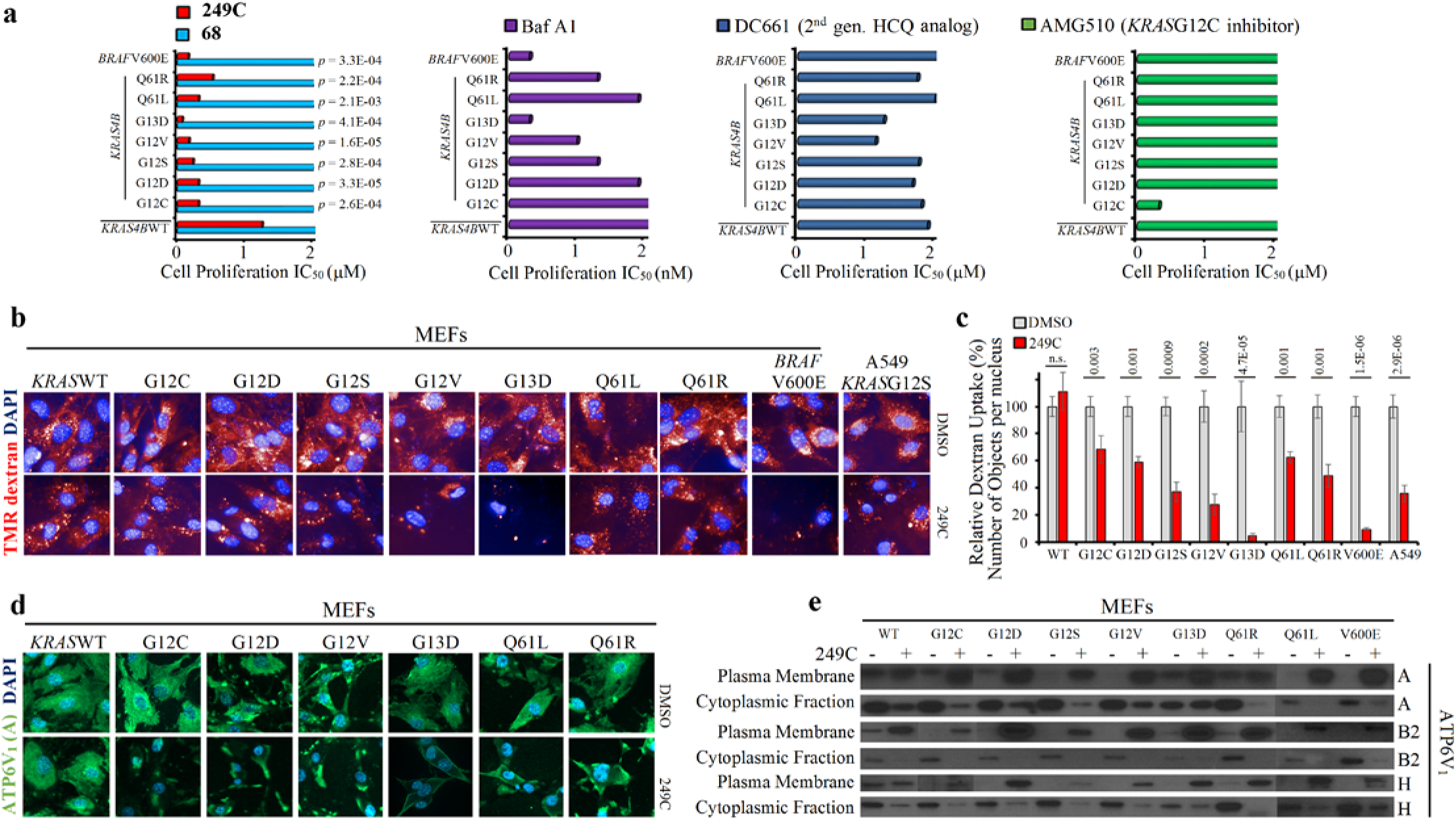
249C Differentially Inhibits Fibroblasts Bearing Mutations in *KRAS* and *BRAF* via Macropinocytosis and Translocation of V-ATPase to the Plasma Membrane. (**A**) Mouse embryonic fibroblasts (MEFs) bearing only single point mutations in human *KRAS/BRAF* treated with 249C. *p* values calculated relative to WT cells treated with 249C. Molecule **68** shown for comparison. G13D (0.07 μM), G12V (0.15 μM), and *BRAF*V600E (0.11 μM) and are exquisitely sensitive to 249C treatment with IC_50_ values about 20-fold and 8-fold lower than WT (1.25 μM). Cell proliferation assays of stable cell lines obtained from the NCI treated withV-ATPase inhibitors 249C and BafA1, HCQ analog (DC661), and *KRAS*G12C inhibitor (AMG510). For each bar, n = 3. Representative of 3 biological replicates is presented. (**B**) Effect of V-ATPase inhibition using 249C on macropinocytosis in mutant *KRAS* and *BRAF* MEFs and A549 (*KRAS*G12S) cancer cells. Fluorescence micrographs showing tetramethylrhodamine (TMR)–dextran uptake. (**C**) Quantification of TMR– dextran uptake. The number of objects per nucleus was quantified using high-content imaging Harmony® PhenoLOGIC™ software in each biological replicate (n = 3); each replicate had 5 DMSO and 5 249C-treated samples; data are mean ± stdv.; two-tailed Student’s t-test. (**D**) Effect of 249C treatment on V-ATPase localization in MEFs. Fluorescence micrographs of MEFs bearing WT and mutant *KRAS* cells immunostained for V1A subunit; >3 images were captured per well. Representative of 2 biological replicates presented. (**E**) Immunoblots of ATP6V1-A, B2, and H in the plasma membrane and cytoplasmic fraction of MEFs. Representative of 2 biological replicates presented

V-ATPase can localize to both the lysosomal and plasma membranes (PM); PM V-ATPase was reported to control MP in RAS mutants^10^ and OligomycinA, an ATP synthase inhibitor, is reported to alter K-Ras PM localization^31^. We next tested whether 249C-mediated inhibition of V-ATPase might impact Ras/Raf-induced MP in a mutant-specific manner. Indeed, treatment with 249C caused the strongest inhibition of MP in MEFs bearing *KRAS*G13D (94%) and *BRAF*V600E mutations (90%), in agreement with the low IC_50_ of cells bearing these mutants (**Fig. 5b-c**). We next investigated the effect of 249C treatment on the subcellular distribution of V-ATPase in these MEFs. Upon 249C treatment, whereas in MEFs harbouring WT *KRAS*, localization of V-ATPase was predominantly cytoplasmic, the presence of *KRAS*G13D led to pronounced accumulation of V-ATPase at the PM, as determined by immunofluorescence using a ATP6V1A subunit-specific antibody (**Fig. 5d**), and substantiated by subcellular fractionation using V1A, B2 and H antibodies (**Fig. 5e**). These data underscore a link between V-ATPase trafficking and nutrient supply by MP that could be exploited to curb metabolic adaptation vulnerabilities; in particular 249C may robustly inhibit *KRAS*G13D mutant tumors for patient-tailored drugs.

### Pharmacodynamic Safety and Anti-tumor Efficacy of 249C in a Mouse Xenograft Model

Finally, to test whether 249C could inhibit autophagy and attenuate the growth of mutant *KRAS*-dependent cancers as a single agent *in vivo*, we evaluated 249C in a mouse xenograft model of NSCLC. Briefly, we injected mutant *KRAS* A549 cells into the flanks of nude mice to establish xenografts, randomly assigned mice to vehicle control (n = 7) or 249C-recipient groups (n = 7, 10 mg/kg) with twice daily i.p. injections for 3 weeks, and recorded tumor volume and body weight over time. During the course of the study, all mice survived but tumor volume was significantly lower in 249C-treated mice (**Fig. 6a**, *p* = 2.9 × 10^−7^). We subsequently sacrificed all mice and excised their tumors for further investigation. Tumor mass was lower in the 249C-treated group (**Fig. 6b**, *p* = 5.3 × 10^−6^), and tumors from mice treated with 249C showed upregulation of LC3 I/II, as assessed by immunoblotting, indicative of inhibition of autophagic flux (**Fig. 6c**). Pharmacokinetic studies revealed that after 4 hours, a safe maximum concentration (C_max_) of 20 μM of 249C was detected in blood (or plasma) of mice, confirming metabolic stability (**Fig. 6d**). Furthermore, 150 mg/kg of 249C was determined to be the maximum tolerated dose in mice as body weight did not change in either group, suggesting a lack of toxicity from treatment (**Fig. 6e**). To further evaluate the potential for secondary effects and adverse events in human cells, which is key to preventing Phase I failures, we used BioMAP® Phenotypic Safety and Toxicology profiling^32^. In human primary cells relevant to physiology, 249C was determined to have no adverse events at 4 doses across over 100 biomarker readouts, an endorsement of its candidacy to progress into the clinic (**Fig. 6f**). Finally, we assessed the effect of 249C on human hERG. Whereas the autophagy inhibitors CQ and HCQ are known to induce a high risk of cardiac electrocardiogram Long QT Syndrome (LQTS) through inhibition of hERG, 249C showed no risk at 3.7 μM (**Fig. 6g**) with an IC_50_ of 14.4 μM (data not shown), whereas HCQ showed an IC_50_ of 5 μM, indicating a sizeable therapeutic window^18.33^. Collectively, these data indicate that 249C is efficacious in inhibiting tumor growth with minimal side effects in our investigations thus far, establishing *in vivo* proof-of-concept for targeting autophagy/MP with 249C in mutant *KRAS* cancers.

**Figure 6:**
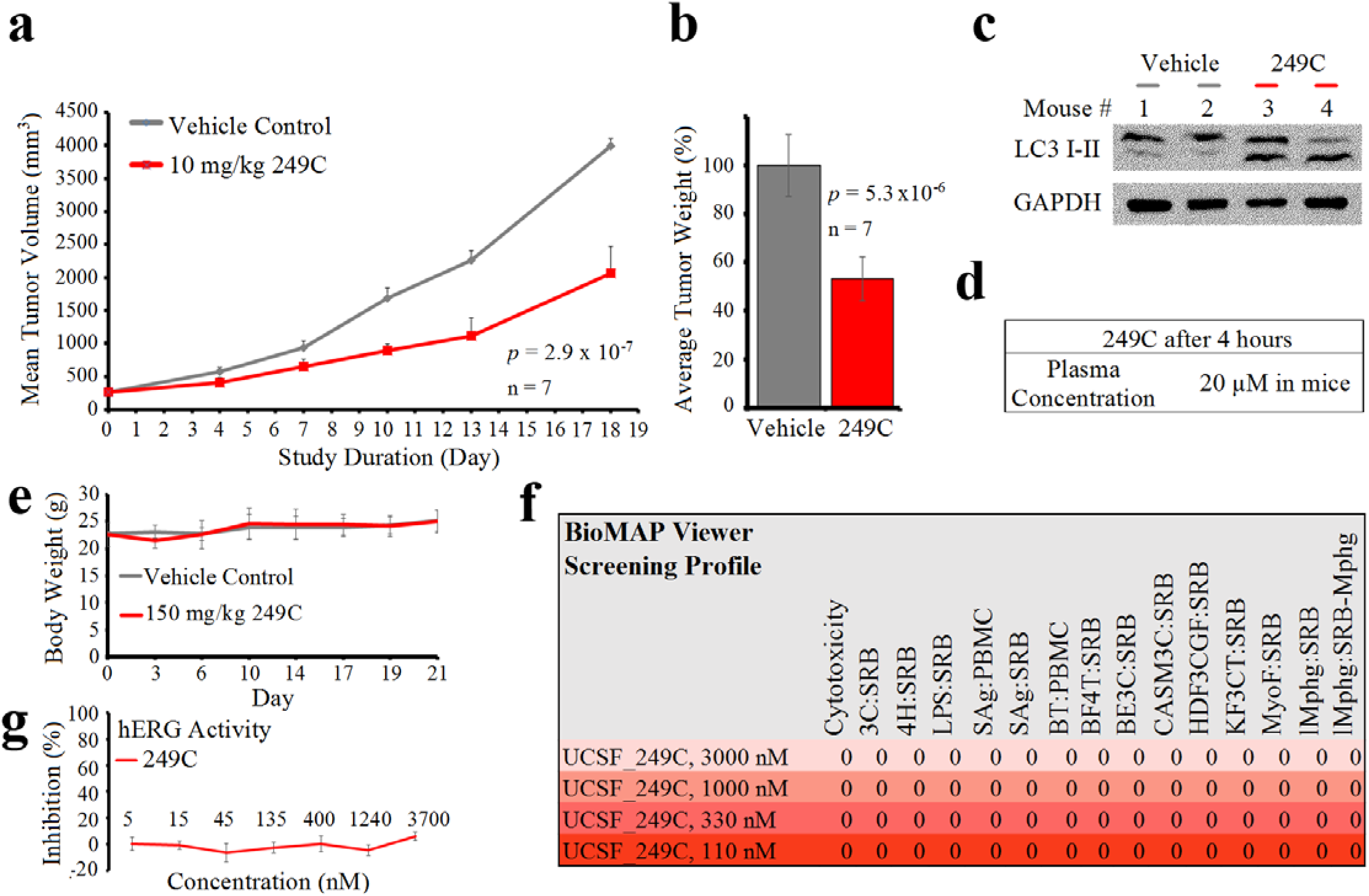
249C Treatment Inhibits *in vivo* Growth of Mutant *KRAS* NSCLC in a Mouse Xenograft Model. (**A**) 5-week-old female athymic mice were inoculated with 5 × 10^6^ NSCLC A549 cells in the lower flank region to establish tumors and randomly assigned to treatment regimens of DMSO vehicle control or 249C (10 mg/kg × twice a day, i.p.). Changes in tumor volume of mice treated with the vehicle control or 249C (n = 7, *p* = 2.9 × 10^−7^) over the course of the study are shown. (**B**) At the end point, tumor mass was determined on harvested tumors by weighing (n =7, *p* = 5.3 × 10^−6^). (**C**) *Ex vivo* analyses of 249C-treated tumors show an up-regulation of LC3-I/II expression when compared with the DMSO controls in lung cancer xenografts. (**D**) After 4 hours, a safe maximum concentration (C_max_) of 20 μM of 249C was detected in blood (or plasma) of mice. (**E**) The safest highest dose of 249C measured is 150 mg/kg as no significant difference in body weight was observed after 21 days *in vivo*. (**F**) *in vitro* BioMap® Safety and Toxicology screening profile of 249C reveals no adverse events at 4 doses across over 100 biomarker readsouts (see methods for additional details). (**G**) Percent inhibition in fluorescence polarization assays of human *ether-à-go-go*-related gene (hERG) responsible for electrical activity of the heart by 249C is close to 0 up to doses of 3,700 nM. Data are mean ± stdv.; two-tailed Student’s t-test for all graphs presented.

## DISCUSSION

In this study, we describe the identification of 249C, a lead compound for treatment of *KRAS* mutant cancers that kills cancer cells by targeting a druggable hotspot within the V-ATPase subunit ATP6V1H and thereby inhibiting autophagy/MP. The mechanism of 249C is distinct from those of other V-ATPase inhibitors such as BafA1 and ConcanamycinA (antifungals), which target ATP6VoC, as well as other autophagy inhibitors such as HCQ and CQ (antimalarials), which act through a different target, PPT1. While both antifungals and antimalarials have potential therapeutic activity, their clinical utility is hampered by toxicity. BafA1 and Con-A have been relegated as laboratory compounds, and HCQ and CQ show substaintial hERG activity. By contrast, 249C shows minimal cytototoxicity compared to BafA1 and minimal inhibition of hERG while maintaining nanomolar potency.

Identification of the relevant mechanisms of action and cellular targets has been a critical barrier to exploiting small molecules with therapeutic potential from being developed into approved drugs for patient benefit. Here, three independent assays – chemo-proteomics, comparative profiling, and CRISPR-based chemical-genetic screens – pointed to V-ATPase as the molecular target of 249C, which we then confirmed using cell-based, biochemical and biophysical techniques. Compellingly, 249C inhibited proton pumping and ATP hydrolysis through direct binding to V-ATPase. ATP6V1H is part of the stator complex^34^, and is suggested to regulate bridging of rotor (Vo) and stator (V_1_) complexes and physically preventing ATP-driven rotation; upon H subunit binding, 249C could conceivably alter V_1_-Vo bridging and hamper biochemical activity. The physical coupling model proposed here is in alignment with previous reports where blocking V-ATPase using BafA1/Con-A results in lysosomal inhibition^35,36^. The identification of 249C’s molecular target now facilitates rational, structure- and target-guided engineering and selection of targeted patient groups and treatment applications. Indeed, in a xenograft model of mutant *KRAS* NSCLC, 249C shows promising *in vivo* activity, low toxicity, and favorable pharmacokinetics relative to FDA-approved drugs.

While the emergence of the covalent *KRAS*G12C inhibitor being developed by Amgen has garnered enthusiasm, G12C accounts for only 11% of *KRAS* mutations across all human cancers. Further, mutations in codon 13 confer worse prognosis and outcomes than those in codon 12^37,38^ underscoring the need for new target-specific therapeutic strategies. In characterizing the ability of 249C to inhibit cell proliferation, autophagy, and MP in cell lines with different *KRAS*/*BRAF* mutations, we found that potency depends on mutant context; G13D showed the highest sensitivity. Thus, V-ATPase inhibition may impact individual *KRAS/BRAF* mutants uniquely, matching findings that have identified signaling differences between individual Ras mutations; G13D is reported to exhibit differential GTP-hydrolysis^39^ and could differentially modulate mRNA levels of V-ATPase subunits like G12D does^40^. Interestingly, K-Ras4b has been shown to interact with ATP6V0A2^41^ and it’s conceivable that individual Ras mutants could show differential interactomes. Thus, there may not be a single anti-Ras therapeutic approach for all RAS mutant cancers; instead, mutation-specific therapeutic strategies must be deployed for different RAS mutations. Selective targeting of PM V-ATPase could offer a strategy for the development of therapeutics against specific *RAS* mutant tumors emphasizing an opportunity to identify patient cohorts that might most benefit from treatment. More broadly, autophagy dysregulation has been linked to a wide range of diseases including metabolic disorders, aging, and neurodegenerative diseases^42^, thus motivating investigation of 249C as a drug-candidate for other treatment applications.

## METHODS

### Cell lines, Reagents and Proliferation Assays

Most cells were obtained from American Type Culture Collection (ATCC) and were cultured in ATCC-recommended media supplemented with 10% fetal bovine serum and 2% antibiotics (Penicillin-Streptomycin). MDA-MB-231 CRISPRi cells were a gift from Hani Goodarzi at UCSF. MEFs expressing human *KRAS/BRAF* were from the Frederick National Laboratory for Cancer Research, The National Cancer Institute. Baf A1, Chloroquine, Hydroxychloroquine, and DC661 were from Medchemexpress LLC. Rapamycin was from Selleckchem and Torin was from Fisher. DNA transfection into cells was performed with TransIT®-LT1 Transfection Reagent (Mirus Bio) by following the manufacturer’s recommendations.

Logarithmically growing cells were plated in antibiotic-free medium supplemented with 2% fetal bovine serum at a density of 5,000 cells per well. The next day, cells were treated in triplicates with increasing doses of in-house small molecule inhibitor drugs and a DMSO vehicle control for 3 days and subsequently assessed for cell viability by measurement of ATP with CellTiter-Glo Luminescent Cell Viability Assay (Promega). Signal intensity was read on a Glomax™ 96 Microplate Luminometer (Promega) and percent cell survival was calculated based on the reading of vehicle control cells set as 100%. Each compound was tested a minimum of 2 times with n = 3.

### Chemical Synthesis of 249C and Structure-Activity Relationship (SAR) Investigation

249C and other small molecules (~300) were synthesized at UCSF. Briefly, rationally designed small molecules to represent diversity in the compound library were synthesized. Compounds synthesized in this series are potent in multiple cancer cell lines. Most calculated properties meet Lipinski’s RO5 (drug-likeness) criteria.

### Mutation Analysis in CCLE Cell Lines

The Cancer Cell Line Encyclopedia (CCLE, www.broadinstitute.org/ccle) was used to download somatic mutation data for over 1000 cancer cell lines^1^. We filtered the data to retain only mutations (single nucleotide variants, SNVs, and insertion deletions, indels) detected in 53 cell lines that were included in our MTS assays (Fig. 1). Non-coding variants, silent mutations and in-frame insertions were removed, as these were not considered functional. We retained in-frame deletions as these may remove essential binding sites. The Combined Annotation Dependent Database (https://cadd.gs.washington.edu/) was used to assess the predicted functional importance of SNVs, and we retained only variants with CADD Phred scores ≥15^2^. Including these predicted damaging SNVs as well as indels (minus in-frame insertions), we determined the number of times each gene was mutated across the 53 cell lines. Genes with multiple mutations in the same cell line were only counted once. Multiple Wilcoxon rank sum tests were carried out, using the “lapply” looping function in the R statistical environment, to test for association between presence/absence of mutations in each gene and IC_50_ values in each cell line. A Wilcoxon rank sum test was also carried out for Ras/Raf mutations, which included combined mutation data for genes *KRAS*, *NRAS*, *HRAS*, and *BRAF*.

### Mass Spectrometric Whole Proteome Analysis

After treatment with small molecules, A549 cells were trypsinized, lysed in 8 M Urea, 1% SDS, 50 mM Tris pH 8.5, protease (Complete) and phosphatase (PhoStop) inhibitors, and samples were sent to The Thermo Fisher Center for Multiplexed Proteomics to be processed as described^3^.

### BioMAP and Benchmarking

BioMAP assays comprising 12 human primary cell-based systems (co-cultured cells simulating 12 different disease contexts) were treated in the presence or absence of test agents (known drugs and our 249C compound) as described^7^. All cells were from a pool of multiple donors (n = 2 – 6) and collected in accordance with appropriate regulatory protocols. Direct ELISA was used to measure selected biomarker level readouts (adhesion receptors, cytokines, enzymes etc.) and activity profiles (normalized data sets) were generated for each test agent. The resulting profiles from biomarker readouts were compared or ‘benchmarked’ to a known 4000-molecule database by statistical methods to identify similarities and mechanistic insights described previously^8^ to provide Pearson and Z scores.

### Immunoblotting

Protein extracts were prepared from drug-treated cells by lysis on ice for 20 minutes in M-PER Mammalian Protein Extraction Reagent (ThermoScientific) supplemented with protease and phosphatase inhibitors (Roche). Lysates were centrifuged at 16,000 RPM for 20 minutes at 4°C and the supernatants were assessed for concentration via a Bradford assay. 25-50 μg of protein extracts were used for gel electrophoresis followed by immunoblotting onto PVDF membranes. Blots were probed with the following primary antibodies: GAPDH (ThermoFisher), SQSTM1 and LC3 (Cell Signaling).

### Electron Microscopy

A549 cells attached to ACLAR film^9^ were treated for 24 hours with 249C (250 nM) or Baf A1 (10 nM) or DMSO as a control. The cells on film were then fixed by immersion in 2% glutaradehyde in 0.08 M Na-cacodylate buffer, pH 7.3 containing 2 mM CaCl_2_ that had been prewarmed to 37 ° C for 1 hour with gentle agitation during which the fixative cooled to room temperature. The samples were then rinsed with 0.1 M Na-cacodylate buffer at room temp., post-fixed with 1% OsO_4_ containing 1.5% potassium ferrocyanide in 0.1 M Na-cacodylate buffer for 45 min on ice, rinsed with water, *en bloc* stained with 3.5% uranyl acetate in water for 1 hour at room temp., dehydrated in ethanol followed by propylene oxide, and embedded in Eponate 12 resin (Ted Pell, Inc, Redding, CA). Thin sections were cut with a Leica UCT ultramicrotome using a Diatome diamond knife and picked up on Pioloform films on slot grids. The grids were then post-stained with 1% uranyl acetate followed by Sato’s lead^10^. Sections were imaged with an FEI T12 TEM equipped with a Gatan U895 4k × 4k camera at 120 kV. Quantification was performed with custom Python image analysis software.

### Lysosome Staining

Cells were treated with the respective molecules in 2-well chambered coverglass slides at a final concentration of 1 μM for 18 hours followed by addition of lysosome-specific dyes. LysoTracker™ Red DND-99 and pH sensitive-LysoSensor™ Yellow/Blue DND-160 purchased from ThermoFisher were added to the cells for 1 hour at 37° C along with Hoescht to stain DNA according to the manufacturer’s recommendations. Live cell images were captured at 20X with a Zeiss spinning disc confocal & TIRF Fura microscope at UCSF’s Laboratory for Cell Analysis.

### CRISPR Screen

Genome-scale screens using MDA-MB-231 CRISPRi cells (a gift from Hani Goodarzi at UCSF) were performed in a similar manner as previously described^11,12^. For genome-wide knockdown, CRISPRi-v2 sgRNA libraries^12^ [targeting 18,905 genes and marked with Blue Fluorescent Protein (BFP)] were transduced into MDA-MB-231 CRISPRi cells at an MOI < 1 (percentage of transduced cells 2 days after transduction: ~30%). Cells were maintained in DMEM in 20 × T-182.5 flasks for the course of the screen. After transduction, the cells were selected with puromycin for 2 days, at which point transduced cells accounted for 90% of the population assessed for BFP by flow cytometry. Following 1 day of recovery without puromycin, samples at time-point t_0_ with a minimum established coverage of >1,000 cells/sgRNA were harvested and the remaining cells were split into two populations for untreated growth (DMSO control) and 249C-treated growth. The cells were maintained in T-182.5 flasks at an average coverage of greater than 1000 cells per sgRNA for the duration of the screen. For 249C treatment, 525 nM 249C was added to the cells at Day 0 and Day 5 and removed the following day. Cells were harvested on days 19 (DMSO) and 21 (249C) (6.1 doubling differences between treated and untreated populations). Genomic DNA was isolated and the sgRNA-encoding region was amplified and processed for next generation sequencing on an Illumina HiSeq 4000 as described previously^13^. Sequencing reads were aligned to the CRISPRi library sequences, counted, and quantified using the Python-based ScreenProcessing pipeline available at https://github.com/mhorlbeck/ScreenProcessing^12^. Generation of negative control genes and calculation of phenotypes and Mann-Whitney p-values was performed as described previously^11^. Sensitivity phenotypes (ρ) were calculated by calculating the log2 change in enrichment of an sgRNA in the treated and untreated samples, subtracting the equivalent median value for all non-targeting sgRNAs, and dividing by the number of population doubling differences between the treated and untreated populations^11,13,14^. Similarly, untreated growth phenotypes (γ) were calculated from the untreated and t_0_ samples, dividing by the total number of doublings of the untreated population. Phenotypes from sgRNAs targeting the same gene were collapsed into a single sensitivity phenotype for each gene using the average of the top three scoring sgRNAs (by absolute value) and assigned a p-value using the Mann-Whitney test of all sgRNAs targeting the same gene compared to the non-targeting controls. For genes with multiple independent transcription start sites (TSSs) targeted by the sgRNA libraries, phenotypes and p-values were calculated independently for each TSS and then collapsed to a single score by selecting the TSS with the lowest Mann-Whitney p-value. Read counts and phenotypes for individual sgRNAs are available in Table SI3. Gene-level phenotypes are available in Table SI4. All additional analyses were performed using python3.6 using a combination of Numpy (v1.15.4), Pandas (v0.25.3), and Scipy (v1.4.1).

### Individual CRISPR Re-tests

Oligonucleotides used to construct sgRNA plasmids directed at the selected genes for individual re-tests were acquired from IDT and cloned into the vector backbone followed by DNA transformation, extraction, and sequencing to verify accuracy and packaging of lentivirus. After 5 days, transduced CRISPRi cells were left untreated (DMSO) or treated with 525 nM of 249C or 30 nM BafA1 for 24 h and assessed for blue fluorescent protein (BFP) using an LSR II flow cytometer. Experiments were performed in triplicates. To validate screening results, we measured the effects of knocking down all 9 subunits on 249C sensitivity in individual re-tests. We infected MDA-MB-231 CRISPRi cells with constructs expressing sgRNAs targeting V-ATPase subunits (*ATP6V1 – A, B2, C1, D, H, F* and *ATP6V0 – A2, C, E1*) or a non-targeting control sgRNA (*Gal4-4*) and used flow cytometry to monitor how the fraction of sgRNA-expressing cells changed after treatment with 249C. MDA-MB-231 CRISPRi cells expressing ATP6V1H subunit targeting sgRNA were depleted more compared to others (Fig. 3c), thus reproducibly indicating that *ATP6V1H* levels affect 249C sensitivity. Thus, 249C’s pattern of drug-gene interactions closely resembles those of V-ATPase-inhibiting agents, particularly that of Baf A1, which binds to the ATP6VoC subunit.

### Subcellular Fractionation

Cytoplasmic and membrane extracts from A549 cells left untreated (DMSO) or treated with 249C (20 μM) or Baf A1 (1.5 μM) for 2 hours were prepared using a Subcellular Protein Fractionation Kit by following the manufacturer’s recommendations (Fisher). Fractions were subjected to Western blotting as described above and blotted for ATP6V1B2 (a gift from Dennis Brown at Harvard University) and ATP6VoD (Abcam) antibodies.

### Protein Purification

Mammalian V-ATPase complex: the entire V-ATPase complex (800 kDa) was purified as described previously from porcine kidneys^23^ using the detergent glycol-diosgenin.

ATP6V1H (H-subunit): Addgene Plasmid #14658 was was expressed in Rosetta2(DE3)pLysS cells and induced with IPTG. Harvested lysates were loaded onto a GSTrap column and bound protein eluted with reduced glutathione. Thrombin was added to cleave the GST tag followed by laoding onto a BioRad SEC650 size-exclusion column.

### Bio-Layer Interferometry Studies for Binding Affinity Determination

Biotinylated-249C was diluted in PBS + 0.05% Tween + 0.2% BSA, pH 7.4 and loaded onto Octet Streptavidin (SA) Biosensors (ForteBio) by following the manufacturer’s recommendations on an Octet RED384 machine (ForteBio, PALL Octet System). Biotin-249C immobilization was checked via Octet® Software prior to introduction of the entire V-ATpase complex and the individual H subunit protein diluted in blocking buffer (1X PBS + 0.05% Tween 20 + 0.2 % BSA + 10 μM Biotin) to eliminate non-specific binding. Graphical output from the Octet® Software of representative data from two independent experiments is presented.

### Proton Pumping Assay in Mammalian Cells

HEK 293T cells were either left untreated (DMSO control) or treated with 10.5 μM 249C or 0.5 μM Baf A1 for 1 hour at 37 °C. This assay was performed as described previously^15^.

### Proton Pumping Assay in Yeast

Spheroplasts from wild-type yeast (a gift from the Walter lab at UCSF) were left untreated (DMSO control) or treated with 29 μM 249C, or 1 μM Baf A1 for 2.5 hours with shaking at 30 °C and vacuolar membrane vesicles were isolated as previously described^20^. Proton transport was measured using ATP-dependent quenching of 9-amino-6-chloro-2-methoxy-acridine (Acridine Orange, ThermoFisher) fluorescence quenching for isolated vacuoles as previously described^21^

### Biochemical V-ATPase Assay

V-ATPase was purified as described previously from yeast^22^ using the detergent dodecylmaltoside and porcine kidney^23^ using the detergent glycol-diosgenin. ATPase assays with purified V-ATPase were performed as described previously. 249C (mammalian: 0.1, 1, 10, 100 μM and yeast: 5, 10 and 100 μM) or Baf A1 (1 μM) were added to the reaction and compared with a negative control to which only DMSO was added. Enzyme-coupled ATPase biochemical activity assays were performed in a 96-well plate as described previously^24^.

### Macropinosome Visualization and Quantification

Macropinosome visualization was essentially done as previously described^29^. Images were captured using an Operetta CLS™ fluorescent microscope for High-Content Analysis (PerkinElmer). Image analysis and quantification was performed with a spot segmentation algorithm using the Perkin Elmer Harmony® PhenoLOGIC™ 3.2 software. Briefly, the images were segmented and individual nuclei (DAPI stained) and cell bodies (by digital phase contrast imaging) were identified. The TMR signal was used for the spot segmentation analysis block. Number of spots, area, and TMR fluorescence intensity were calculated for each cell region, and displayed as mean values per cell. Data are presented as spots - number of objects per nucleus. For each cell line, 5 DMSO control and 5 249C-treated wells were imaged and a minimum of 3 images were captured per well. Data representing three independent experiments are presented.

### Immunofluoresence Assays

Cells were seeded onto glass coverslips and subsequently serum-starved for 24 h. After serum starvation, cells were treated with 249C for ~15 hours and fixed with 4% formaldehyde for 30 min at room temperature. The following sequential steps took place at room temperature: cells were washed twice with PBS, permeabilized (0.1% Tween in PBS) for 10 min, and blocked (5% bovine serum in PBS) for 30 min. The following primary antibody was used: ATP6V1A (Abnova). A corresponding alexafluor-488 secondary antibody was used (ThermoFisher). Cells were mounted onto slides using Mounting Media (Vector Laboratories Inc) containing DAPI. A minimum of 3 images per well were captured using a spinning disk confocal microscope (Zeiss). Data representing two independent experiments are presented.

### Plasma Membrane Fractionation

The plasma membrane protein extraction kit (BioVision) was used to separate the plasma membrane fraction from other cellular membranes according to the manufacturer’s recommended protocol. Data representing two independent experiments are presented.

### Animal Studies, Plasma Concentration, Body Weight

5 × 10^6^ A549 cells were subcutaneously injected into the lower left flank of 6-8-week old female athymic mice (Charles River). To assess establishment of tumors, mice were examined 5 days post-inoculation and were randomly segregated into four treatment groups [vehicle control, 10 mg/kg 249C IP, n = 7 per group] and were treated twice daily with the 249C or vehicle control. Tumor volumes were calculated using the formula (L × W × H)/2 with recorded caliper measurements. Survival curves and statistical analyses were performed using Excel. All animal procedures were performed under IACUC approved protocols and guidelines. Animals from both groups were weighed every 3 days. Plasma concentration was measured from blood collected using standard and established protocols.

### Ex vivo Analysis

At the end point, tumors were harvested, weighed and minced up. Total protein from frozen tissues was prepared with T-PER (ThermoFisher Scientific) supplemented with protease and phophatase inhibitors (Roche). Equal quantities of proteins were combined with 5X protein loading buffer and separated by SDS-PAGE followed by PVDF membrane transfer. Membranes were blocked with 5% milk followed by incubation with LC3 (Cell Signaling) and GAPDH (ThermoFisher) antibodies. Blots were developed with ECL Reagents (Pierce).

### hERG assay

The fluorescence polarization-based assay to characterize the affinity of 249C for human *ether-a-go-go*-related gene (hERG) responsible for electrical activity of the heart was performed as previously described^30^.

### BioMAP

The BioMAP Toxicity Screening Profile was performed as described previously^31^.

### Statistical analyses

All experiments were repeated a minimum of two times with duplicate or triplicate samples. Values including controls are expressed as the mean ± SD. Two-tailed paired Student’s t test was used to test for differences between two groups unless indicated otherwise. Differences with a p≤0.05 were considered as statistically significant.

Correspondence and requests for materials should be addressed to B.T.

## AUTHOR CONTRIBUTIONS

**Conception and design:** B.T., M.J., J.S.W. **Funding support and acquisition:** B.T., J.S.W. **Preparation of the manuscript:** B.T., M.J., J.S.W. All other authors provided feedback. **Analyses and interpretation of data:** B.T., A.C., Y.Z.T., R.F., A.J.S., T.V., J.L.R., M.J., J.S.W.; M.J. analyzed and interpreted CRISPRi data. **Acquisition of data:** B.T. screened ~300 molecules via cellular assays, calculated IC_50_ values, screened 249C in 53 cancer cell lines, prepared samples for whole proteome mass spec, comparative profiling, performed immunoblotting, performed LysoSensor and LysoTracker experiments. B.T. performed the CRISPRi screen and individual re-tests with guidance from M.J. and J.S.W. B.T. performed subcellular fractionation, mammalian proton pumping assays, yeast proton pumping assays, assisted with H subunit protein purification, and performed Octet/BLI binding studies. B.T. performed cellular assays in MEFs, MP assays with dextran, immunofluorescence staining, plasma membrane isolation, and *ex vivo* analysis. A.C. captured microscopy images and assisted in MP high content analysis. Y.Y. and I.B.S. performed and supported chemical synthesis, respectively. Y.Z.T. purified mammalian V-ATPase used for Octet/BLI studies, performed mammalian V-ATPase assays, and analyzed the data. R.F. performed electron microscopy and analyzed data. A.J.S. performed statistical analysis for WT and mutant IC_50_ values using CCLE. T.V. performed yeast V-ATPase assays and analyzed data. C.R.L. isolated DNA for next-gen sequencing for CRISPRi. M.J. provided extensive technical expertise on CRISPRi screening. B.T. performed/oversaw the remainder of assays.

## ACKNOWLEDGMENTS

This study was supported in part by UCSF’s Helen Diller Family Comprehensive Cancer Center (HDFCCC) Laboratory for Cell Analysis Shared Resource Facility through a grant from NIH (P30CA082103. Flow Cytometry was performed at the UCSF Core facility (RRID: SCR_018206) that is supported in part by the NIH DRC Center Grant (P30 DK063720). The authors thank the following individuals for their generous contributions: Dr. Hani Goodarzi at UCSF (MDA-MB-231 CRISPRi cells) and Dr. Dennis Brown at Harvard University (ATP6V1A and ATP6V1B2 antibodies used for Figure 4). The MEF cells expressing human *KRAS* or *BRAF* were from the Frederick National Laboratory for Cancer Research, National Cancer Institute. BT thanks Drs. Keith Yamamoto and Jaynta Debnath from UCSF for valuable discussions and career guidance. We thank Drs. Michael Forgac and Stephan Wilkens for technical discussions. We thank Morgan Boone from the Walter lab at UCSF for yeast and related reagents. We thank Thoracic Oncology for space and assistance with chemical synthesis/reagents. Figures 3 and 4 were created with BioRender.com.

## REFERENCES

1. Simanshu DK, Nissley DV, McCormick F. (2017) RAS Proteins and Their Regulators in Human Disease. Cell 170:17–33.

2. Cox AD, Fesik SW, Kimmelman AC, Luo J, Der CJ. (2014) Drugging the undruggable Ras: mission possible? Nat Rev Drug Discov. 11:828–851.

3. Waters AM, Der CJ. (2018) KRAS: The Critical Driver and Therapeutic Target for Pancreatic Cancer. Cold Spring Harb Perspect Med. 8(9): a031435.

4. Hobbs AG, Der CJ, Rossman KL. (2016) RAS isoforms and mutations in cancer at a glance. J Cell Sci. 129(7): 1287–1292.

5. Choi AM, Ryter SW, Levine B. (2013) Autophagy in human health and disease. N Engl J Med. 368:651–662.

6. Guo JY, Teng X, Laddha SV, Ma S, Van Nostrand SC, Yang Y, Khor S, Chan CS, Rabinowitz JD, White E. (2016) Autophagy provides metabolic substrates to maintain energy charge and nucleotide pools in Ras-driven lung cancer cells. Genes Dev. 30: 1704–17.

7. Efeyan A, Comb WC, Sabatini DM. (2015) Nutrient-sensing mechanisms and pathways. Nature 517(7534):302–10.

8. Mizushima N, Levine B, Cuervo AM, Klionsky DJ (2008) Autophagy fights disease through cellular self-digestion. Nature 451;1069–75.

9. Guo JY, White E. (2013) Autophagy is required for mitochondrial function, lipid metabolism, growth, and fate of KRAS(G12D)-driven lung tumors. Autophagy 9:1636–8.

10. Ramirez C, Hauser AD, Vucic EA, Bar-Sagi D. Plasma membrane V-ATPase controls oncogenic RAS-induced macropinocytosis. Nature. 2019;576(7787):477–481.

11. Rabinowitz, J.D. & White, E. Autophagy and metabolism. Science 2010;330:1344–1348.

12. Commisso, C., Davidson, S.M., Soydaner-Azeloglu, R.G., Parker, S.J., Kamphorst, J.J., Hackett, S., et al. Macropinocytosis of protein is an amino acid supply route in Ras-transformed cells. Nature 2013;497:633–637.

13. Forgac, M. Vacuolar ATPases: rotary proton pumps in physiology and pathophysiology. Nat. Rev. Mol. Cell Biol. 8, 917–929 (2007).

14. Zhao J, Benlekbir S, Rubinstein JL. Electron cryomicroscopy observation of rotational states in a eukaryotic V-ATPase. Nature. 2015 May 14;521(7551):241–5.

15. Stransky L, Cotter K & Forgac M The function of v-ATPases in cancer. Physiol. Rev 96, 1071–1091 (2016).

16. Li Z, Du L, Zhang W, et al. Complete elucidation of the late steps of bafilomycin biosynthesis in *Streptomyces lohii*. J Biol Chem. 2017;292(17):7095–7104.

17. Rebecca VW, Nicastri MC, Fennelly C, et al. PPT1 Promotes Tumor Growth and Is the Molecular Target of Chloroquine Derivatives in Cancer. Cancer Discov. 2019;9(2):220–229.

18. Michaud V, Dow P, Al Rihani SB, Deodhar M, Arwood M, Cicali B, Turgeon J. Risk Assessment of Drug-Induced Long QT Syndrome for Some COVID-19 Repurposed Drugs. Clin Transl Sci. 2020 Sep 5.

19. Rebecca, V, Amaravadi, R. Emerging strategies to effectively target autophagy in cancer. Oncogene 35, 1–11 (2016).

20. Carew, J. S. and Nawrocki, S. T. (2017). Drain the lysosome: Development of the novel orally available autophagy inhibitor ROC-325. Autophagy.

21. McAlister GC, et al. MultiNotch MS3 enables accurate, sensitive, and multiplexed detection of differential expression across cancer cell line proteomes. Anal Chem. 2014; 86:7150–7158.

22. Berg EL, Kunkel EJ, Hytopoulos E, Plavec I. Characterization of compound mechanisms and secondary activities by BioMAP analysis. J Pharmacol Toxicol Methods. 2006;53(1):67–74.

23. Jost M, Weissman JS. CRISPR Approaches to Small Molecule Target Identification. ACS Chem Biol. 2018 Feb 16;13(2):366–375.

24. Gilbert LA, Larson MH, Morsut L, Liu Z, Brar GA, Torres SE, Stern-Ginossar N, Brandman O, Whitehead EH, Doudna JA, Lim WA, Weissman JS, Qi LS. CRISPR-mediated modular RNA-guided regulation of transcription in eukaryotes. Cell. 2013 Jul 18;154(2):442–51.

25. Horlbeck MA, Gilbert LA, Villalta JE, Adamson B, Pak RA, Chen Y, Fields AP, Park CY, Corn JE, Kampmann M, Weissman JS. Compact and highly active next-generation libraries for CRISPR-mediated gene repression and activation. Elife. 2016 Sep 23;5:e19760.

26. Gilbert LA, Horlbeck MA, Adamson B, Villalta JE, Chen Y, Whitehead EH, Guimaraes C, Panning B, Ploegh HL, Bassik MC, Qi LS, Kampmann M, Weissman, JS. (2014) Genome-Scale CRISPR-Mediated Control of Gene Repression and Activation. Cell 159: 647–61.

27. Bowman BJ, Bowman EJ (2002) Mutations in subunit C of the vacuolar ATPase confer resistance to bafilomycin and identify a conserved antibiotic binding site. JBiolChem 277: 3965–3972.

28. Liberman, R., Bond, S., Shainheit, M. G., Stadecker, M. J., and Forgac, M. (2014) Regulated assembly of the V-ATPase is increased during cluster disruption-induced maturation of dendritic cells through a PI-3 kinase/ mTOR-dependent pathway. J. Biol. Chem. 289, 1355–1363

29. Stolze B, Reinhart S, Bulllinger L, Fröhling S, Scholl C. (2015) Comparative analysis of KRAS codon 12, 13, 18, 61, and 117 mutations using human MCF10A isogenic cell lines. Sci Rep. 5:8535.

30. Drosten M, Dhawahir A, Sum EY, Urosevic J, Lechuga CG, Esteban LM, Castellano E, Guerra C, Santos E, Barbacid M. (2010) Genetic analysis of Ras signaling pathways in cell proliferation, migration and survival. EMBO J. 29:1091–104.

31. Salim AA, Tan L, Huang XC, et al. Oligomycins as inhibitors of K-Ras plasma membrane localisation. Org Biomol Chem. 2016;14(2):711–715.

32. Kleinstreuer NC, Yang J, Berg EL, et al. Phenotypic screening of the ToxCast chemical library to classify toxic and therapeutic mechanisms. Nat Biotechnol. 2014;32(6):583–591.

33. Trudeau MC, Warmke JW, Ganetzky B, Robertson GA. HERG, a human inward rectifier in the voltage-gated potassium channel family. Science. 1995;269(5220):92–5.

34. Wilkens S, Inoue T, Forgac M. Three-dimensional structure of the vacuolar ATPase. Localization of subunit H by difference imaging and chemical cross-linking. J Biol Chem. 2004;279:41942–41949.

35. Zoncu R et al. mTORC1 senses lysosomal amino acids through an inside-out mechanism that requires the vacuolar H(+)-ATPase. Science 334, 678–683 (2011).

36. Bar-Peled L, Schweitzer LD, Zoncu R & Sabatini DM Ragulator is a GEF for the rag GTPases that signal amino acid levels to mTORC1. Cell 150, 1196–1208 (2012).

37. Tze-Kiong Er, Chih-Chieh Chen, Luis Bujanda, Marta Herreros-Villanueva, Clinical relevance of KRAS mutations in codon 13: Where are we? Cancer Letters, 343;1, 2014, 1-5.

38. Hobbs GA, Der CJ. Binge Drinking: Macropinocytosis Promotes Tumorigenic Growth of RAS-Mutant Cancers. Trends Biochem Sci. 2020 Jun;45(6):459–461.

39. Rabara D, Tran TH, Dharmaiah S, Stephens RM, McCormick F, Simanshu DK, Holderfield M. KRAS G13D sensitivity to neurofibromin-mediated GTP hydrolysis. Proc Natl Acad Sci U S A. 2019 Oct 29;116(44):22122–22131.

40. Yang J, Guo F, Yuan L, Lv G, Gong J, Chen J. Elevated expression of the V-ATPase D2 subunit triggers increased energy metabolite levels in Kras^G12D^ -driven cancer cells. J Cell Biochem. 2019 Feb 11.

41. Zhang X, Cao J, Miller SP, Jing H, Lin H. Comparative Nucleotide-Dependent Interactome Analysis Reveals Shared and Differential Properties of KRas4a and KRas4b. ACS Cent Sci. 2018 Jan 24;4(1):71–80.

42. Rubinsztein, D. C., Codogno, P. & Levine, B. Autophagy modulation as a potential therapeutic target for diverse diseases. Nat. Rev. Drug Discov. 11, 709–730 (2012).

